# CUL5^WSB2^ uses BCL2 proteins as co-receptors to target BIM for degradation

**DOI:** 10.1101/2025.08.14.670414

**Authors:** Wilhelm Vaysse-Zinkhöfer, Enya Marie Catherine Alcindor, Nicholas Garaffo, David Paul Toczyski

## Abstract

Anti-apoptotic BCL2 family proteins (e.g. BCL-XL) protect cells by binding and inhibiting pro-apoptotic proteins (e.g. BIM). Although CUL5^WSB2^ was linked to apoptosis regulation, its substrate and mechanism were unknown. We find that BCL2 proteins recruit BIM to CUL5^WSB2^ for degradation. WSB2 recognizes BCL-XL through a motif conserved between BCL-XL, BCL-W and BCL2, but not MCL1. Disruption of this interaction through mutation of either BCL-XL or WSB2 blocks the binding of WSB2 to the BCL-XL/BIM dimer. WSB2 also associates with the MCL1/BIM dimer through a separate WSB2 interface, suggesting that WSB2 has evolved independent two means to target BIM. While WSB2 is not essential in most cells, it is essential in cells derived from tumors of the nervous system, and knockdown of WSB2 in these lines causes death and apoptosis. This work uncovers a novel mechanism of apoptosis regulation, with implications for developing therapies against neuroblastomas and other cancers reliant on this pathway for survival.

## Introduction

The apoptosis pathway triggers cell death in response to either extracellular or intracellular signals^1^. Apoptosis is mediated by BAX, BOK and BAK, which oligomerize to form pores in the outer mitochondrial membrane through their four BH domains^1–3^. Antagonizing this event are a set of anti-apoptotic proteins (BCL2, BCL-XL, BCL-W, MCL1, and Bfl-1/A1) which use these same domains to bind pore-forming proteins and block their oligomerization^1,4^. This event is further regulated by two sets of proteins that contain only one BH domain, BH3. One set of these “BH3-only” proteins (e.g. BID, BIM and PUMA), also known as “activators”, binds and activates the pore formers directly, and also binds and interferes with anti-apoptotic BCL2-like proteins^5–9^. The second set of BH3-only proteins (e.g. NOXA, BAD and BIK) sequester the anti-apoptotic BCL2-like proteins, and thus activate apoptosis by inhibiting its inhibitors^1,5,9,10^.

This fine balance of pro and anti-apoptotic factors is further regulated in myriad ways. BIM, a pro-apoptotic factor mutated in non-small cell lung cancer (NSCLC)^11–13^, Leukemia^14,15^ and Neuroblastomas^16^, is regulated at the level of splicing, phosphorylation, localization and protein turnover^17^. BIM exists in three main splicing isoforms: extra-long (BIM_EL_), long (BIM_L_) and short (BIM_S_)^17,18^. Amino acids encoded in exons specific to BIM_EL_ are phosphorylated by both ERK1/2 and RSK1/2^19^ (Fig. 2A). These phosphorylations promote the targeting of BIM_EL_ by the F-box protein ßTRCP, which acts as a substrate specificity factor for the SKP1-CUL1-F-Box complex (SCF)^19^. The SCF is one of seven Cullin-Ring Ligases (CRLs), each of which contains a different Cullin (CUL1 for the SCF) and one of two RING subunits (Rbx1 or Rbx2)^20–22^. The human genome encodes several hundred genes with domain structures that correspond to CRL substrate receptors, most of which are thought to be genuine CRL subunits^23^. CUL5 employs Rbx2, and two adaptors, ELOB and ELOC, which tether any of about 40 SOCS-box proteins to CUL5^24–26^. Each of the CRL ligases works in concert with a separate ubiquitin ligase that adds the first ubiquitin to the substrate. Most CRLs use Arih1 for this function, whereas the CUL5 ligase uses a paralog, Arih2^27–29^. Ubiquitin ligases are essential in a number of pathways involved in development, cell division, specification or migration^30^ and are often mutated or amplified in tumors^31–34^. Consequently, significant efforts have been made to more readily identify their targets^35–38^.

Anti-apoptotic BCL2 family members such as BCL2, BCL-XL, and BCL-W have long been known to suppress apoptosis by binding, and thereby inhibiting, the function of pro-apoptotic BH3-only proteins^1,7,39,40^. We have found that BCL-XL inhibits at least one pro-apoptotic BH3-only protein, BIM, by facilitating its degradation. Here, we show that the CUL5 substrate adaptor WSB2 binds BCL-XL directly and, through that interaction, targets the associated BIM for degradation without affecting the half-life of BCL-XL itself. While WSB2 is not essential in most primary or tumor cells, high-throughput data from *The Cancer Map Dependency Project* (DepMap) suggest it is critical for survival in tumors of the peripheral nervous system, such as neuroblastoma. We show loss of WSB2 in neuroblastoma cells leads to a sudden and dramatic increase in apoptosis, suggesting that WSB2 is a strong candidate for drug discovery in this class of cancers.

## Results

### WSB2 and CUL5 mutants have high levels of BIM_EL_, BIM_L_ and BIM_S_ splicing isoforms

Previously, high throughput studies showed that mutation of CUL5, RBX2, ARIH2, or WSB2 sensitized HAP1 cells to the mitochondrial complex II inhibitor TTFA^41^, and WSB2-depleted HAP1 cells displayed a growth defect after TTFA treatment (Fig. 1A). Additionally, analysis of dependency data from the DepMap consortium revealed that WSB2 positively correlated with anti-apoptotic BCL-W, BCL2, MCL1, BCL-XL (Pearson’s corr > 0.2) and negatively correlated with pro-apoptotic PMAIP (NOXA) and BAX (Pearson’s corr < 0.2) (Fig. 1B), consistent with data from several CRISPR screens^42,43^. GO analysis of the top 100 WSB2 co-dependencies revealed significant enrichment in apoptosis regulation (Fig. 1B). As these results suggested a potential role for WSB2 in negative apoptosis regulation, we investigated the potential involvement of WSB2 in this pathway.

**Figure 1:**
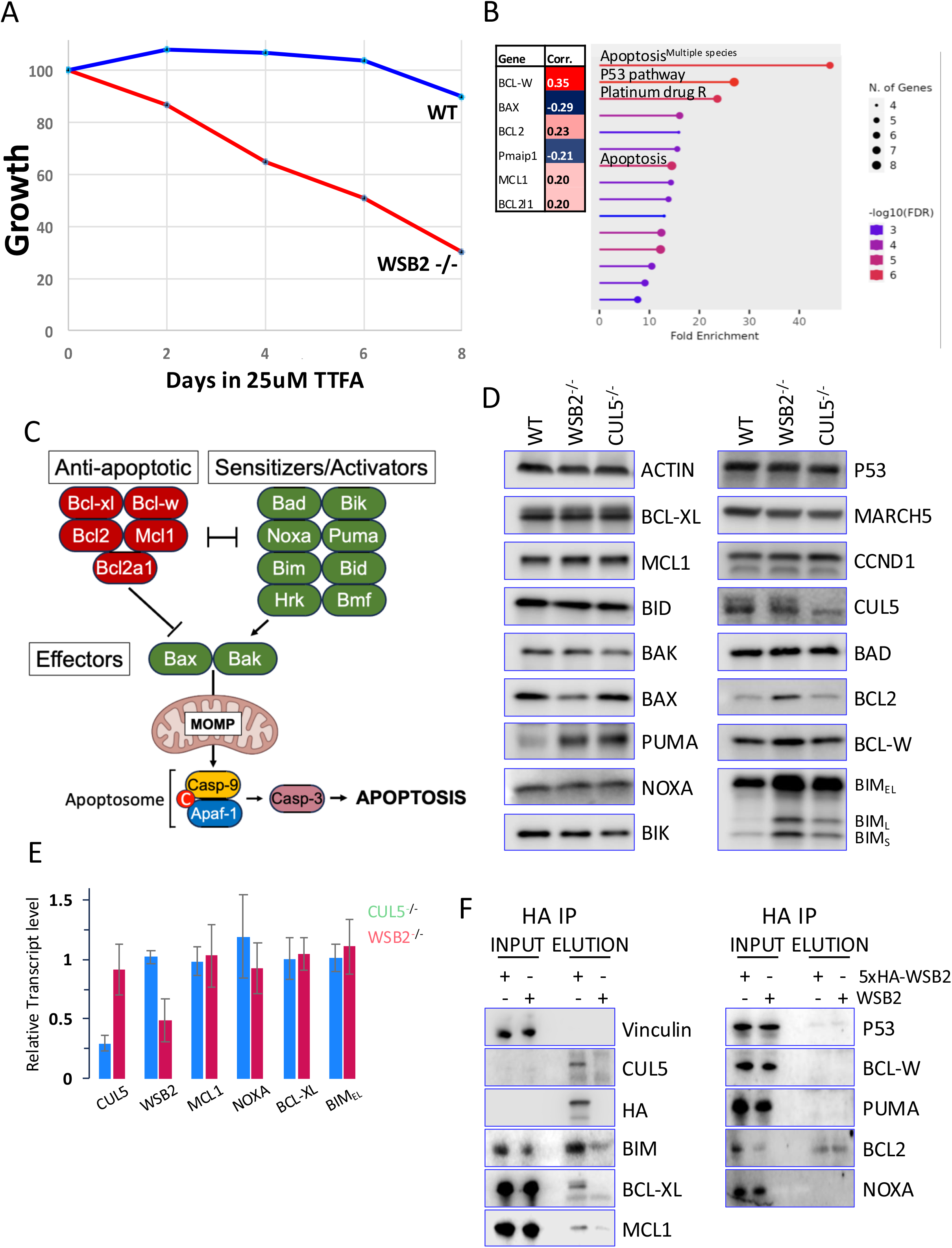
WSB2 and CUL5 mutants have high levels of BIM_EL_, BIM_L_ and BIM_S_. **A.** Competitive growth assay comparing cells expressing sgRNAs against WSB2 (red line) or control sgRNAs (blue line) in Cas9 positive HAP1 cells in the presence of 25 µM TTFA, over an 8-day period. **B**. Co-dependencies data from the DepMap consortium and Gene Ontology (GO) classification of the top 100 WSB2’s co-dependencies. **C**. Schematic representation of the intrinsic apoptotic pathway. **D**. Immunoblot analysis of apoptotic proteins in clonal WSB2 or CUL5 mutant RPE1 cells. Quantitation (Fig. S1B). **E**. qPCR analysis of transcript levels in WSB2 or CUL5 mutant RPE1s cells normalized to WT. **F**. 5xHA-WSB2 or an untagged WSB2 allele was induced with dox overnight and samples were immunoprecipitated with anti-HA antibodies.

Interactions between members of the BCL2 family are thought to dictate the inhibition or initiation of apoptosis. The BH3-only sensitizers/activators (PUMA, BIK, BAD, HRK, BMF and NOXA) are upregulated through p53 and MYC in response to apoptotic stimuli such as DNA damage or mitochondrial stress^44,45^. These proteins subsequently bind the anti-apoptotic BCL2 proteins (BCL2, BCL-W, BCL-XL, and MCL1) freeing the BH3-only activators BIM and BID and the effectors BAX and BAK, which are usually sequestered by the anti-apoptotic members. The release of BIM and BID leads to mitochondrial outer membrane permeabilization (MOMP) through the activation of BAX and BAK, committing cells to apoptosis^3,4^ (Fig. 1C).

To test WSB2’s function in this process, we generated clonal RPE1 cell lines with WSB2 or CUL5 mutations, confirmed by sequencing (Fig. S1A) and, for CUL5, by immunoblotting (Fig. 1D). Commercially available antibodies directed against WSB2 were not functional, making direct detection challenging. To assess the impact of WSB2 or CUL5 mutation on apoptotic protein levels, we probed immunoblots for a large panel of apoptotic proteins (PUMA, NOXA, BIK, BAD, BCL2, BCL-W, BCL-XL, MCL1, BID, BIM, BAK and BAX) as well as p53, MARCH5, CUL5 and Cyclin-D1 (CCND1) (Fig. 1D, S1B).

Immunoblotting analysis revealed a significant upregulation of all BIM isoforms across multiple independently derived WSB2 mutants and a significant upregulation of BIM_L_ across multiple independently derived CUL5 mutants. BCL2 protein levels showed an upward trend but were not significantly different from WT (Fig. S1B). We also observed a modest increase in PUMA levels, although the signal varied considerably between experiments. In contrast, the steady state levels of BIK, BAD, BCL-W, BCL-XL, MCL1, BID, BAK and BAX remained unchanged in the WSB2 and CUL5 mutants (Fig. 1D, S1B). Moreover, the levels of MARCH5 (a ubiquitin ligase important for apoptosis, whose mutation also rendered cells sensitive to TTFA^41^) and p53 were also unaffected. Although CCND1 has been suggested to be a potential substrate of WSB2^46^, no changes in CCND1 levels were observed.

Treatment with 1uM Rotenone (mitochondrial complex I inhibitor) or 25nM Camptothecin (CPT; Topoisomerase I inhibitor) during a 48h time course lead to an increased percentage of PARP in the cleaved form (c-PARP) in WSB2 mutants compared to WT, while CPT treatment showed no effect (Fig. S1C), suggesting that WSB2 mutant cells are sensitized to apoptosis induced by mitochondrial poisons. qPCR analysis showed reduced WSB2 and CUL5 transcripts, consistent with nonsense-mediated decay^47^ (Fig. 1E). However, BIM mRNA levels were unchanged, suggesting it is post-transcriptionally regulated. PUMA and BCL2 mRNA levels were too low to quantify.

We then examined whether BIM or PUMA were direct targets of WSB2. We generated RPE1 cells expressing either untagged or 5xHA-tagged WSB2 alleles under the control of a doxycycline(dox)-inducible promoter and induced expression for 8 hours. Immunoprecipitation with anti-HA antibody showed a robust co-immunoprecipitation of HA-WSB2 with endogenous BCL-XL and BIM_EL_, and, more weakly, MCL1. BIM_L_ and BIM_S_ were not detected, likely due to their low abundance in RPE1 cells (Fig. 1F). Despite publication of WSB2 regulating NOXA^48,49^, WSB2 mutation did not induce a change in NOXA steady state levels and NOXA did not co-immunoprecipitate with WSB2.

### WSB2 regulates BIM through BCL-XL and MCL1

BIM_EL_ is a well-described target of the SCF^ß-TrCP1^ complex, which recognizes and ubiquitinates BIM_EL_ after phosphorylation by RSK1/2 and ERK1/2 (Fig. 2A). In contrast, BIM_L_ and BIM_S_ both lack the phosphorylation sites targeted by RSK1/2 and ERK1/2, preventing them from being recognized and ubiquitinated by SCF^ß-TrCP1^ ^17,19^ (Fig. 2A). To determine if WSB2 is specifically interacting with the BIM_EL_ isoform, we introduced a doxycycline-inducible 3xFLAG-tagged BIM_EL,_ BIM_L_ or BIM_S_ construct into RPE1 cells co-expressing either the 5xHA tagged or untagged WSB2. Following a 3-hour dox treatment, immunoprecipitation with anti-HA antibodies revealed that WSB2 interacts with all BIM isoforms, albeit more weakly with BIM_S_ (Fig. 2B).

**Figure 2:**
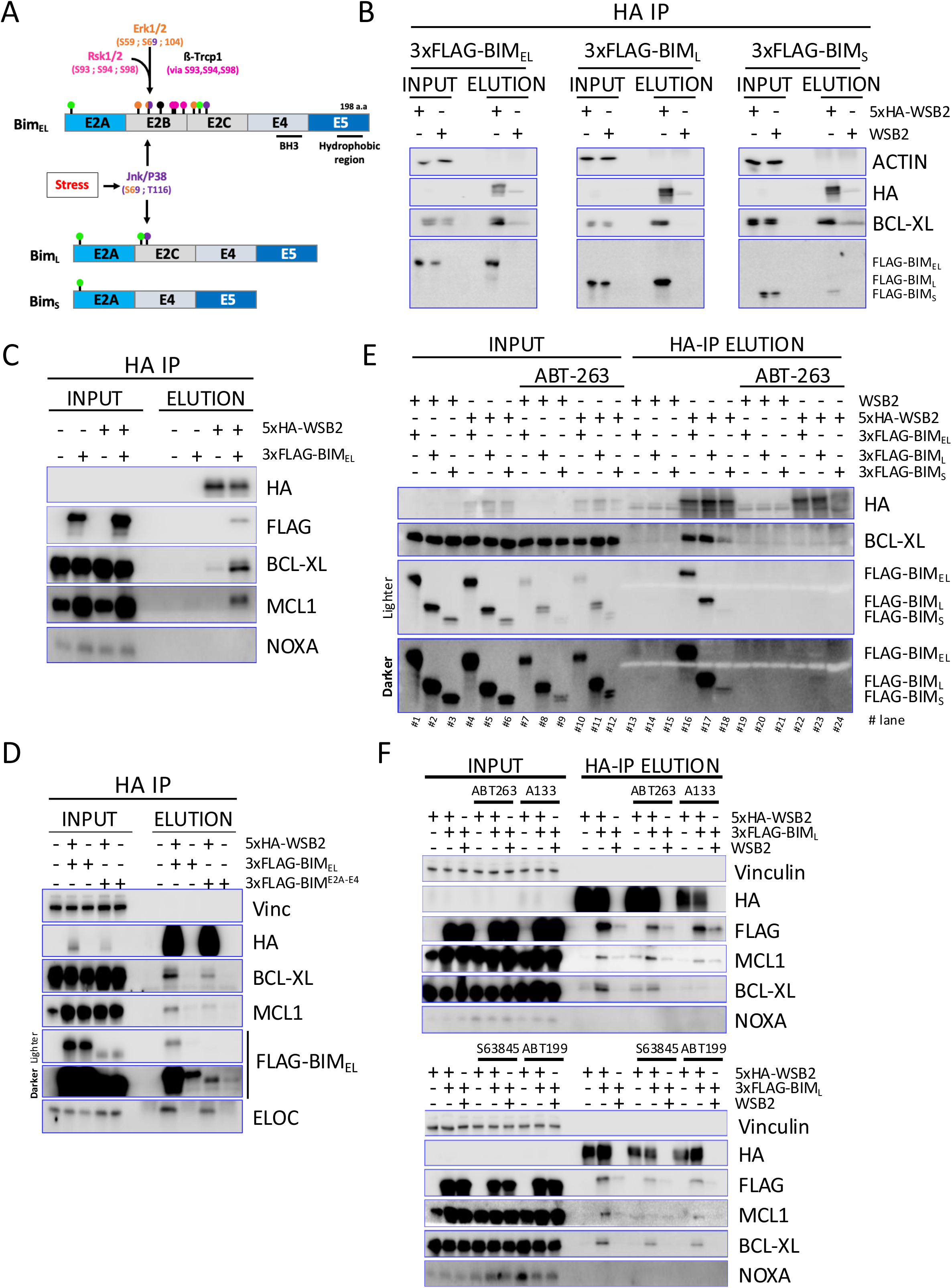
WSB2 regulates BIM through BCL-XL and MCL1. **A**. Schematic representation of the three main BIM splicing isoforms, highlighting their domains and phosphorylation events mediated in part by the MAPK. BIM_EL_ is targeted for degradation by the SCF^βTrCP^ complex following phosphorylation by ERK1/2 and RSK1/2. Green lollipops indicate ubiquitination sites. Pink, orange, and purple lollipops represent RSK1/2, ERK1/2, and JNK/P38 phospho-sites, respectively. **B**. HA-immunoprecipitation of extract from untagged or 5xHA-tagged WSB2 cells co-expressing 3xFLAG tagged BIM_EL_, BIM_L_, or BIM_S_ after 3 hours dox induction. **C**. HA-immunoprecipitation of extracts from cells expressing untagged or 5xHA-tagged WSB2 with or without 3xFLAG-BIM_EL_ co-expression. **D.** HA-immunoprecipitation from cells co-expressing 5xHA-WSB2, 3xFLAG tagged BIM_EL_ or a truncated 3xFLAG-BIM lacking the C-terminal domain (3xFLAG-BIM^E2A-E4^), as indicated.**E**. Same as Figure 2B, except that half the cells were treated with the pan-BH3 mimetic ABT-263 (Navitoclax) for 3 hours. **F**. Co-immunoblot analysis of 3xFLAG-BIM_L_ in the presence of the BH3 mimetics ABT-263, ABT-199, S63845, or A1331852. Immunoprecipitation of 3xFLAG-BIM was used to confirm the efficacy of each BH3 mimetic in disrupting its interaction with the respective anti-apoptotic proteins (Fig. S2C). Dox (to induce 3xFLAG-BIM) and drugs were added for 3 hours before harvesting.

Strikingly, both BCL-XL and MCL1 showed increased interaction with WSB2 upon BIM_EL_ overexpression (Fig. 2C), suggesting WSB2 may specifically recognize a heterodimer. Endogenous BCL-W and BCL2, while detected in whole-cell lysate (Fig. 1D, 1F), were not reproducibly detected in WSB2 HA-IP elutions, possibly due to their low expression in these RPE1 cells (Fig. S2A). The hydrophobic C-terminal E5 domain of BIM is conserved across all three isoforms (Fig. 2A) and is necessary for BIM’s interaction with the mitochondrial outer membrane^50^. We examined whether interaction with the mitochondria was required for BIM’s interaction with WSB2. A 3xFLAG tagged truncated BIM_EL_ lacking the E5 domain (BIM E2A-E4) was overexpressed with or without 5xHA-WSB2. While the BIM E2A-E4 allele was very poorly expressed, it still bound to WSB2, suggesting that the WSB2 BIM interaction is independent of BIM’s interaction with the mitochondrial membrane (Fig. 2D, comparing BIM in lane 9 with background in lane 10). To determine whether WSB2 interacts with BCL-XL and BIM independently, or only within the context of the BCL-XL/BIM heterodimer, we conducted a co-immunoprecipitation experiment in the absence of dimerization by pre-treating cells with the pan-BCL2 BH3-mimetic ABT-263 (Navitoclax) for 3 hours. BH3-mimetics selectively bind to the BH3 domain of anti-apoptotic BCL2-proteins, disrupting their interactions with pro-apoptotic BH3-only proteins, triggering apoptosis. ABT-263 selectively targets BCL-XL, BCL2 and BCL-W. Disruption of the BCL-XL/BIM complex led to a loss of the interaction of WSB2 with BCL-XL and BIM, suggesting that WSB2’s interaction with BIM is largely dependent on BCL-XL (Fig. 2E). To determine whether the loss of interaction between WSB2, BCL-XL and BIM depends on ABT-263 concentration, we performed co-immunoprecipitation experiments using increasing concentrations of ABT-263 (Fig. S2B). Both the WSB2–BCL-XL and WSB2–BIM interactions decreased progressively with loss of co-IP occurring between the 0.4 and 2uM dose. Even cancer cell lines expressing very high levels of BIM have an IC50 for ABT263 of approximately 1uM^51^9/17/26 2:40:00 PM, arguing that disruption of the complex is not an artifact observed only at high concentrations of ABT-263 (Fig. S2B).

To investigate the specificity of these interactions, cells were treated with the BH3 mimetics A1331852, S63845 and ABT199, which target BCL-XL, MCL1 and BCL2, respectively. We carried out this experiment examining BIM_L_ due to its robust binding to WSB2, and to eliminate any complication arising from BIM_EL,_ which is targeted by both CUL5^WSB2^ and SCF^ßTRCP^. Immunoprecipitation of FLAG-BIM_L_ confirmed that each inhibitor effectively disrupted the association between BIM and its respective target proteins (Fig. S2C): ABT-263 reduced BCL-XL, BCL-W (partially), and BCL2, but not MCL1 in the FLAG-IP; A1331852 reduced only BCL-XL; S63845 reduced MCL1 (and BCL2 very slightly); ABT199 reduced only BCL2. Treatment with ABT-263 reduced the interaction between WSB2 with both BCL-XL and BIM_L_, but not with MCL1 (Fig. 2F). Reciprocally, treatment with S63845 disrupted the WSB2-MCL1 interaction, but not the WSB2-BCL-XL interaction. The A1331852 compound primarily disrupted the WSB2-BCL-XL interaction with a slightly less strong effect on BIM_L_. This may be because, in the absence of BCL-XL, BIM is redistributed to other BCL2 family members and is recognized by WSB2 through these interactions. In contrast, ABT199 lead to very little change in the WSB2-BIM_L_ interaction, consistent with its specificity for BCL2 (Fig. 2F). Together, these results suggest that WSB2 primarily targets BIM through the BCL-XL/BIM and MCL1/BIM complexes, but can also recognize BIM in association with other apoptotic proteins.

### WSB2 regulates BIM independently of the ßTRCP1 pathway

To determine whether WSB2 mutation affects apoptotic protein stability, we carried out a cycloheximide time course. BIM_EL_ and BIM_L_ were short-lived, whereas BIM_S_ appears slightly more stable in the WSB2 and CUL5 mutants (Fig. 3A, Fig. S3A, S3B). BCL-XL, BCL2, and NOXA (which undergoes a mobility shift) were stable. Loss of WSB2 stabilized BIM but did not alter BCL-XL or known WSB2 substrates (NOXA, CCND1). Although PUMA and BCL2 levels were slightly elevated in WSB2 mutants (Fig. 1D), their half-lives were not markedly changed (Fig. S3C).

**Figure 3:**
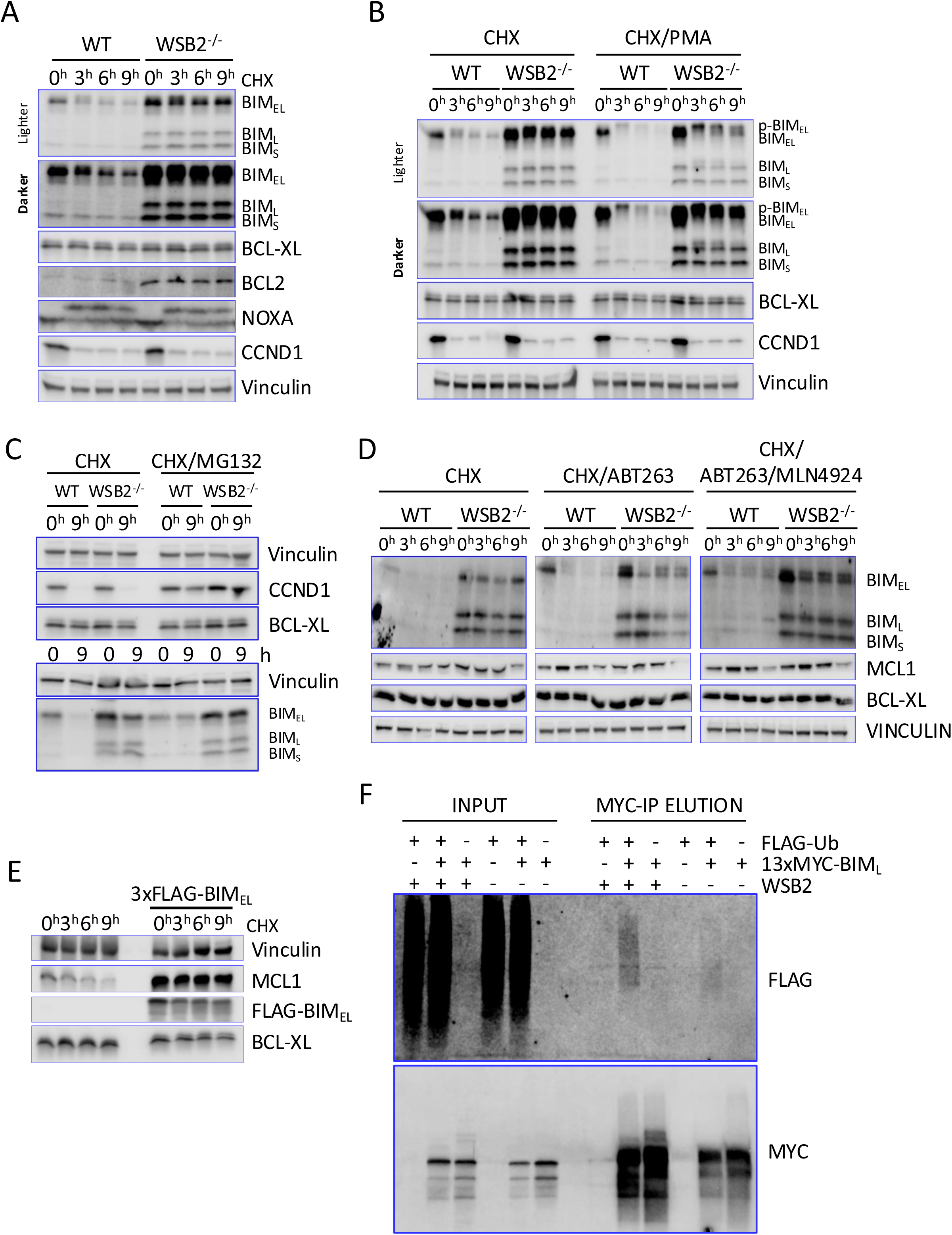
WSB2 regulates BIM independently of the ßTrCP1 pathway. **A**. WT and WSB2 mutant RPE1 cells were treated with cycloheximide and samples were harvested as indicated. Quantitation was normalized to Vinculin loading control and subsequently normalized to the t=0 signal. Phosphoforms of each BIM isoforms were combined for normalization. (Fig. S3A). **B.** As in panel A, except that PMA was added to one set of samples at the same time as cycloheximide. Quantitation Fig. S3D. **C.** As in panel A, except that MG132 was added to one set of samples at the same time as cycloheximide. Quantitation (Fig. S3E). **D.** As in panel A, comparing cells treated with cycloheximide and ABT-263 to cells additionally treated with MLN4924. All drugs were added at t=0. **E**. WT and dox inducible 3xFLAG-BIM_EL_ RPE1 cells were dox induced 3 hours and treated with Z-VAD-FMK (to block apoptosis) before treatment with cycloheximide and samples harvested as indicated. **F**. FLAG-Ubiquitin and 13xMYC-BIM_L_ were co-induced with dox in wild-type or WSB2 mutant RPE1 cells as indicated, and extracts were immunoprecipitated with anti-MYC antibodies.

The longest form of BIM, BIM_EL,_ is regulated through the MAPK pathway. ERK phosphorylates BIM_EL_ serines 59, 69 and 104, which then promotes RSK-mediated phosphorylation of BIM_EL_ on the serines 93, 94 and 98^19^. This enables the recognition and subsequent ubiquitination of BIM_EL_ by the SCF^ßTRCP^ complex. The more potent, smaller isoforms, BIM_L_ and BIM_S,_ both lack the domains required for RSK/ERK phosphorylation and regulation by SCF^ßTRCP1^ (Fig. 2A). Given this, we were surprised to see BIM_EL_ entirely stabilized in the WSB2 mutant. To determine if both CUL5^WSB2^ and SCF^ßTRCP^ target BIM, we performed a time-course experiment following treatment with phorbol 12-myristate 13-acetate (PMA), which has been reported to drive BIM_EL_ destabilization through the RSK/ERK/SCF^ßTRCP1^ pathway^19^. As anticipated, treatment with PMA did not affect the stability of isoforms BIM_L_ and BIM_S_. However, BIM_EL_ was destabilized in the WSB2 mutant when PMA was added (Fig. 3B, Fig. S3D). Moreover, a noticeable phospho-shift was observed in BIM_EL_, consistent with PMA activating ERK/RSK mediated phosphorylation^19^. Interestingly, a phosphoshift of the stable BIM_L_ was also observed. To address whether the destabilization of BIM was dependent on the proteasome, we performed similar time-course experiments treating the cells with 50uM of the proteasome inhibitor MG132 (Fig. 3C, Fig. S3E). Proteasome inhibition stabilizes BIM, suggesting that WSB2 targets BIM for degradation through the proteasome.

To examine the turnover of monomeric BIM, we employed ABT-263. Addition of ABT-263 caused BIM degradation for all three isoforms, even in the WSB2 mutant (Fig. 3D). This is unlikely to be due to SCF^ßTRCP^ since this complex is only known to target the BIM_EL_ spliced form^19^. This suggests that yet another E3 targets monomeric BIM. BIM_EL_ (but not BIM_L_ or BIM_S_) has been reported to be degraded by the proteasome even in the absence of poly-ubiquitination ^52^. However, WSB2-independent turnover in ABT-263 occurred for all three forms of BIM and was reduced by the neddylation inhibitor MLN4924, which inactivates CRL ligases, suggesting that targeting of the monomeric forms of BIM requires, directly or indirectly, a CRL.

Although WSB2 uses BCL-XL to target BIM, BCL-XL itself was stable. It may be that, under normal conditions, only a small fraction of BCL-XL is bound to BIM and targeted by WSB2. To test this hypothesis, we overexpressed BIM_EL_ for 3 hours before treating the cells with cycloheximide. BIM_EL_ overexpression led to only a modest destabilization of BCL-XL in cycloheximide, suggesting that BCL-XL is much less efficiently ubiquitinated (Fig. 3E). Consistent with previous observations (Fig. 2C), BIM_EL_ overexpression strongly increased and stabilized MCL1^53,54^.

To examine whether BIM is ubiquitinated by the CUL5^WSB2^ complex, we introduced a doxycycline-inducible FLAG-Ubiquitin and a 13xMYC-BIM_L_ construct into RPE1 cells. We employed the BIM_L_ isoform because it lacked the confounding effects of being simultaneously targeted by SCF^ßTRCP1^. Cells were treated with 100uM Z-VAD-FMK during the 8h induction with 1ug/ml doxycycline to prevent apoptosis. After harvesting, we performed an immunoprecipitation with anti-MYC antibodies and blotted for FLAG-tagged (Ubiquitin) and MYC-tagged (BIM) proteins. Proteasome inhibitors were not used in these experiments, as we observed that their use led to the disruption of WSB2 and BIM overexpression. Anti-MYC immunoprecipitation resulted in a high-molecular weight FLAG-reactive smear indicative of polyubiquitination only when both FLAG-ubiquitin and 13xMYC-BIM_L_ were co-expressed (Fig. 3F). Mutation of WSB2 resulted in a reduction of the ubiquitin smear associated with MYC pull-down compared to wild-type (Fig. 3F). These results suggest that CUL5^WSB2^ promotes BIM ubiquitination. Residual ubiquitination in the WSB2 mutant may represent a high level of monomeric BIM seen upon overexpression.

### BIM is targeted through a direct WSB2-BCL-XL interaction

Our results indicate that WSB2 regulates BIM exclusively when in complex with another BCL2 family protein. Based on our observation that WSB2 regulates BIM via BCL-XL, we used AlphaFold2 Multimer to predict potential WSB2 interaction interfaces among monomeric BH3-containing apoptotic proteins. AlphaFold^55^ evaluates interface confidence (iPTM), domain positioning (PTM), and structural accuracy (PAE, pLDDT). We considered iPTM > 0.8; PTM > 0.5; PAE < 10 and pLDDT > 77 to be significant. As positive controls, ELOB and ELOC both showed high-confidence binding to WSB2. Among 16 proteins tested, only BCL-XL, BCL2, and BCL-W demonstrated strong interactions with WSB2, whereas the WSB2 paralog WSB1 did not (Fig. 4A; Fig. S4A). These predictions suggested that WSB2 binds BCL-XL via a DKEM motif (D156/K157/E158/M159) in BCL-XL and an ERRRR motif (E28/R157/R174/R297/R353) in WSB2 (Fig. 4B). The DKEM motif is conserved in BCL2 and BCL-W but is absent in MCL1 and BIM_EL_ (Fig. 4C), and no direct WSB2/BIM interaction was predicted (Fig. 4A). *In silico* alanine substitutions in the DKEM to K157A/E158A/M159A (BCL-XL^AAA^) or glutamate substitution in the ERRRR motif to R157E/R174E/R297E (WSB2^EEE^) abolished the predicted interaction (Fig. 4A, Fig. S4B). To experimentally validate our prediction, we generated doxycycline-inducible V5-tagged wild-type and generated these mutant alleles in a 3xFLAG BIM_EL_ background. Despite the fact that the BCL-XL^AAA^ allele expressed better than WT, it had reduced binding to WSB2 (Fig. 4D; Fig. S4C-D). Analogously, WSB2^EEE^ did not interact with BCL-XL, but retained its interaction with MCL1 (Fig. 4E; 4F; 4G; Fig. S4E). Because BCL2 was also predicted to interact with WSB2 through the same motif as BCL-XL, we performed similar co-immunoprecipitation experiments using overexpressed BCL2, as endogenous BCL2 could not be detected. As predicted, WSB2 interacted with BCL2, whereas WSB2^EEE^ had reduced BCL2 binding (Fig. 4G). Together, our results indicate that WSB2 engages BCL-XL and BCL2 through a common interaction interface, whereas its interaction with MCL1 relies on a distinct binding mechanism. However, in the absence of reconstitution experiments using purified proteins, we cannot exclude the possibility that WSB2 makes contacts with the BCL-XL/BIM complex as a whole rather than binding BCL-XL directly and independently of BIM.

**Figure 4:**
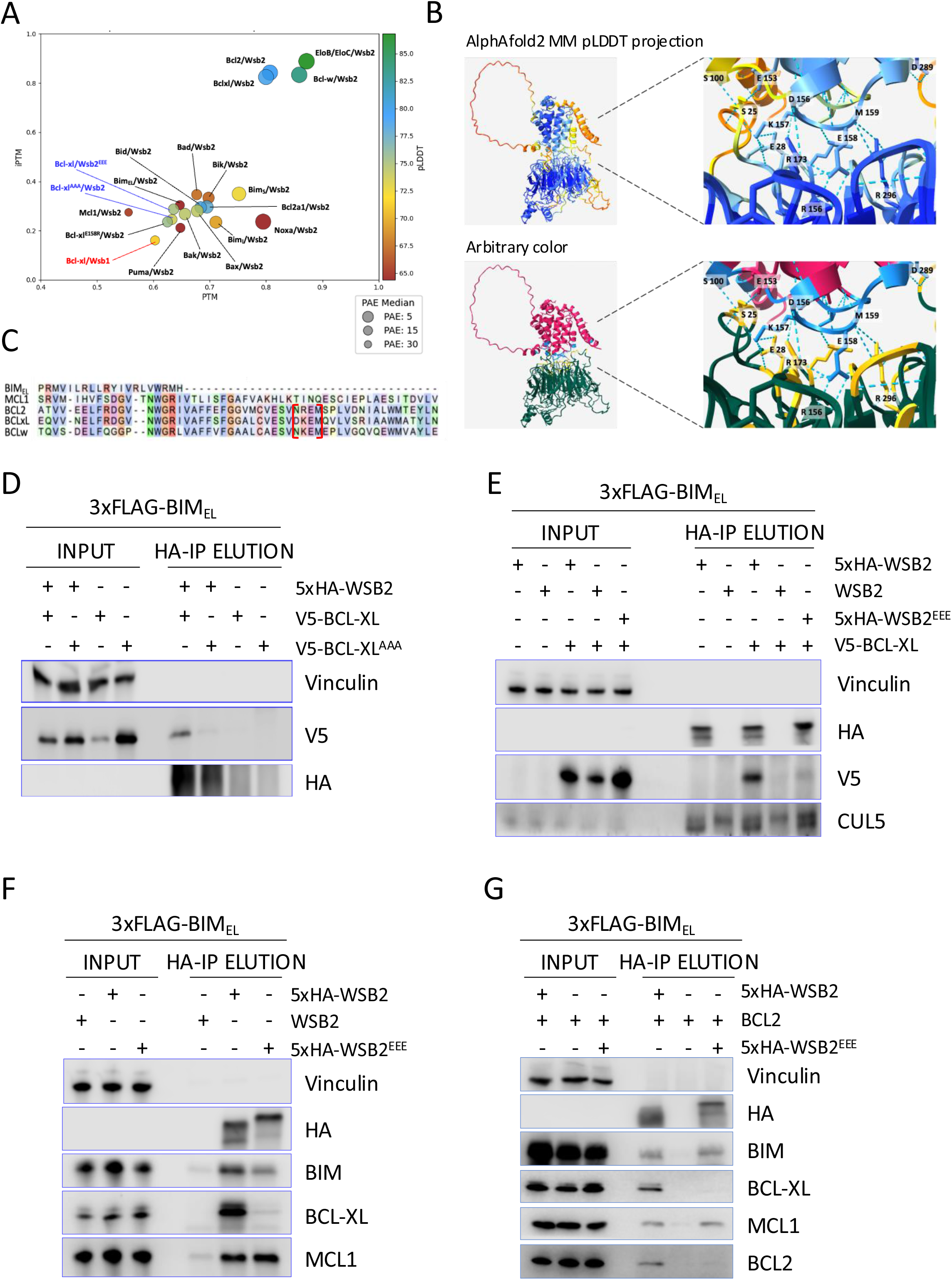
BIM is targeted through WSB2-BCL-XL interaction. **A**. AlphaFold 2 multimer predicted WSB2 dimerization with BCL2 protein family members. The four metrics iPTM (Confidence of protein-protein interaction), PTM (Accuracy of the relative positioning of domains within a multi-domain protein), pLDDT (Confidence in the local structure) and PAE (Expected positional error in the distance between pairs of residues) are shown for dimer prediction between WSB2 and apoptosis proteins. The CUL5 adaptors ELOB and ELOC were used as a positive control. **B**. Top panel represent AlphaFold2 predicted structure of the WSB2/BCL-XL complex with pLDDT coloring. Bottom panel with arbitrary coloring represents AlphaFold2 predicted interaction between WSB2 and BCL-XL through the yellow and light blue residues. **C**. Alignment of BCL-XL, BCL2, BCL-W, MCL1 and BIM showing conservation of the interaction motif between BCL-XL, BCL2 and BCL-W, framed in red. **D**. *In vivo* validation of AlphaFold2 dimer predictions was performed using co-immunoprecipitation assays. A 5xHA-tagged WSB2 allele was co-expressed with either V5-tagged wild-type BCL-XL or BCL-XL^K157A/E158A/M159A^ allele (BCL-XL^AAA^) as indicated. In all cell lines, a 3xFLAG-tagged BIM_EL_ allele was co-expressed. Quantitation (Fig. S4C). **E**. HA-immunoprecipitation in cell lines expressing an untagged, a 5xHA-tagged WT or WSB2^R157E/R174E/R297E^ allele (WSB2^EEE^) and a V5-tagged wild-type BCL-XL, as indicated. In all cell lines, a 3xFLAG-tagged BIM_EL_ allele was co-expressed. **F.** HA-immunoprecipitation in cells expressing an untagged WSB2, 5xHA-tagged WSB2 or a 5xHA-tagged WSB2^R157E/R174E/R297E^ allele. In all cell lines, a 3xFLAG-tagged BIM_EL_ allele was co-expressed. Quantitation (Fig. S4E). **G.** HA-immunoprecipitation in cells expressing an untagged WSB2, 5xHA-tagged WSB2 or 5xHA-tagged WSB2^R157E/R174E/R297E^ allele. In all cell lines, a 3xFLAG-tagged BIM_EL_ allele and BCL2 wild-type allele were co-expressed.

### WSB2 is essential in Neuroblastoma

While WSB2 is expressed in various tissue types, its expression is highest in the nervous system, suggesting a potential role in safeguarding neuronal tissues from apoptosis (Fig. 5A). The Cancer genetic dependencies data from DepMap show that tumors of the nervous system, Neuroblastomas and Malignant Peripheral Nerve Sheath Tumors (MPNSTs), are over-represented amongst those in which WSB2 is essential for viability (Fig. 5B). There is a positive correlation (Pearson Corr. = 0.471) between the essentiality of WSB2 and BCL-W (BCL2L2) in PNS tumors (Fig. 5C). We verified these high through-put studies using a competitive growth assay to measure cell viability. First, we designed constitutively expressed shRNAs against WSB2 and introduced them into the Neuroblastoma cell lines CHP134 and TGW. We employed three shRNAs against WSB2. To verify the efficiency of these shRNAs, we performed qPCR in cells expressing each of the shRNAs (Fig. S5B). The cells were infected with a lentivirus encoding each shRNA and GFP, sorted 24h post-infection and mixed with an equal number of GFP negative cells.

**Figure 5:**
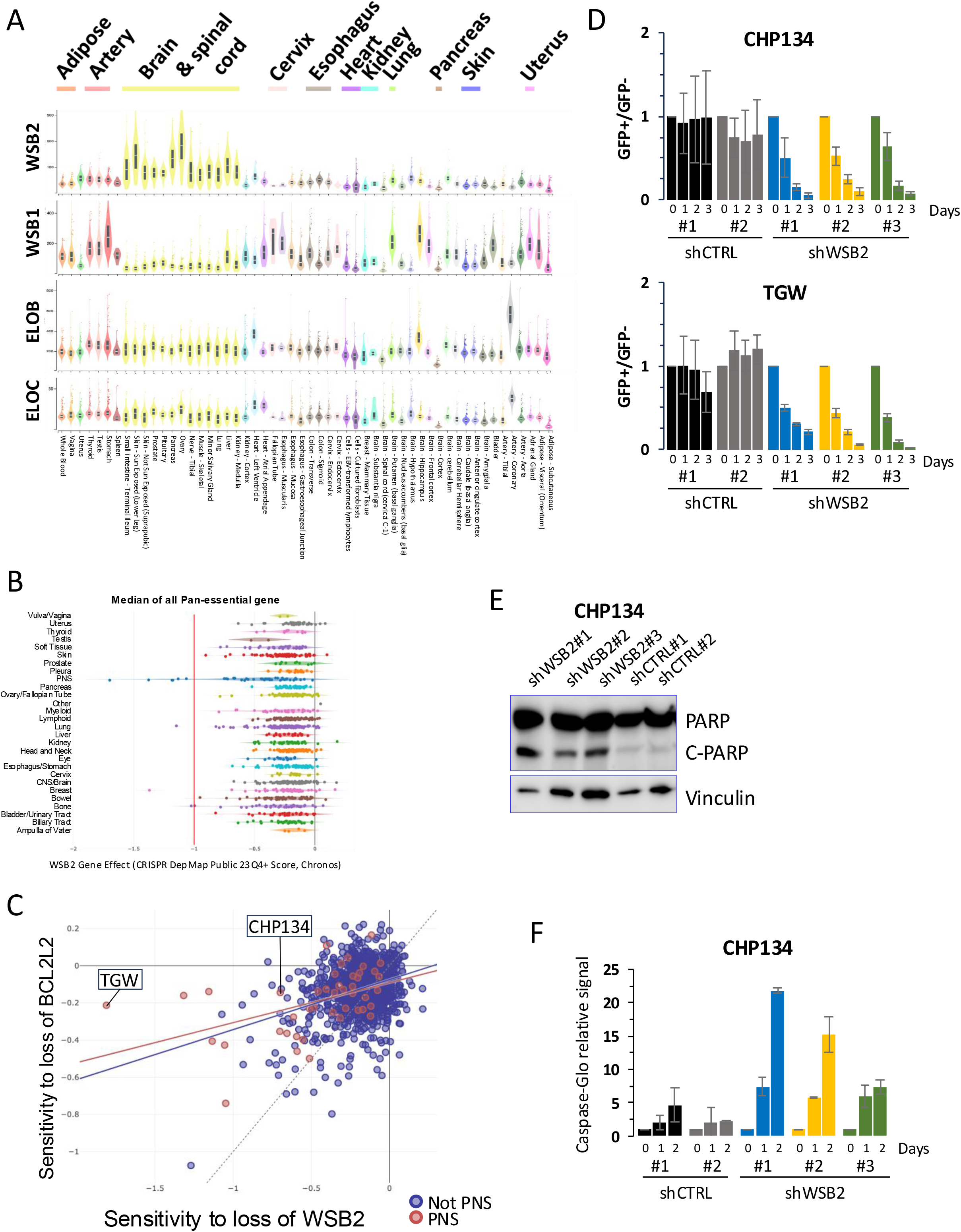
WSB2 is essential in Neuroblastoma. **A.** GTEx portal analysis of transcript levels in bulk tissue RNA sequencing for WSB2, WSB1, ELOB, and ELOC. **B**. DepMap data of tumor sensitivity to WSB2 depletion across multiple cancer types. **C**. DepMap data correlation of tumor sensitivity to WSB2 and BCL2L2 depletion. **D**. Competitive growth assay of CHP134 or TGW cells expressing shRNA against WSB2. (GFP-) cells were mixed in equal number with cells expressing either a shRNA scramble control (GFP+) or a shRNA targeting WSB2 (GFP+). The ratio of GFPpositive to GFP negative cells was measured over 3 days and normalized to t=0. Data are represented as mean ± SD **E**. Immunoblot analysis in CHP134 cells expressing WSB2 shRNA, showing that WSB2 depletion leads to increased levels of cleaved PARP (c-PARP). **F**. Promega Caspase-Glo assay luminescence signal normalized to uninfected control, after transfection with shRNAs against WSB2 or Control over a 2-day time course. Data are represented as mean ± Range.

The ratio of GFP-negative to GFP-positive cells was determined over a 3-day time course using FACS analysis. There was no significant shift in the ratio of cells expressing either control shRNA to the population of GFP-negative cells. In contrast, GFP-positive cells expressing shRNAs against WSB2 were reduced in the population over time (Fig. 5D). To examine CHP134 cells lacking WSB2 in isolation, we sorted GFP-positive cells 24h after infecting with each shRNA encoding virus and assessed the level of cleaved-PARP by western blot (Fig. 5E). We observed a clear induction of c-PARP, compared to total PARP, in cells expressing an WSB2 shRNA, while very little c-PARP could be observed in the control. To further examine this, we used the Caspase-glo assay in which a luminescent signal is emitted proportionally to caspase activity. Cells infected with shRNAs against WSB2 shows clear increased signal compared to control after normalization against non-infected cells (Fig. 5F). Taken together, our data show that loss of WSB2 causes cell death and initiates apoptosis in Neuroblastomas.

## Discussion

A recent CRISPR screen found that CUL5 loss sensitized cells to inhibitors of MCL1 and increased BIM and NOXA levels at both the protein and mRNA levels^56^. Additionally, findings from Hundley et al. showed that mutations of ARIH2, CUL5, RBX2 or WSB2, sensitized HAP1 cells to the mitochondrial complex II inhibitor TTFA^41^. Moreover, high-throughput screening suggested synthetic lethality between WSB2 mutation and MCL1, MARCH5, and BCL-XL^42,43^. These studies suggest that loss of the CUL5^WSB2^ complex somehow primed cells for apoptosis, although it remained unclear if this was a direct effect.

Our study reveals that mutation of CUL5 or WSB2 stabilizes BIM_EL,_ BIM_L_ and, less dramatically, the more stable BIM_S_ isoform (Fig. 3A, B, C and S3A). Notably, WSB2 recognizes these proteins through their association with the stable BCL-XL and MCL1 subunits. This redefines BCL-XL’s and MCL1’s long-known roles in BIM inhibition, showing that they function not only through sequestration, but also as a co-receptor with WSB2 to target BIM. This mechanism operates alongside the well-characterized SCF^ßTRCP1^ pathway and reveals a broader regulatory role for anti-apoptotic BCL2 family members as substrate co-receptors for E3 ligases^19^. Finally, we demonstrate that WSB2 is essential in Neuroblastoma cell lines and protects cells from apoptosis induction, highlighting its potential as a therapeutic target.

Several E3 ligases have been implicated in apoptosis regulation^57,58^. BIM is a known target of SCF^ßTRCP^, and was also suggested to be regulated by Trim2 and CRL2^CIS^ ^59,60^. Similarly, other apoptotic proteins such as BAX, BCL-XL or NOXA have been proposed to be regulated by IBRDC2, RNF183, and CUL5 respectively^61–64^. The pro-apoptotic MCL1 was suggested to be targeted by MARCH5/MULE; SCF^ß-TRCP^,SCF^FBW7^ and TRIM17^34,65–67^. However, in RPE1 cells, we find that BCL-XL, BCL2, NOXA and, to a lesser degree, MCL1 are all stable (Fig. 3), suggesting that, unlike BIM, there may be little bulk turnover of these proteins in unperturbed normal cells. The CUL5^WSB2-BCL-XL/MCL1^ regulation of BIM is reminiscent of the observation that MARCH5 ubiquitinates MCL1 exclusively when NOXA and MCL1 are in complex^65^, showing the generality of BCL2 family members acting as E3 ligases co-receptors.

Although WSB2 depletion did not induce apoptosis in RPE1 cells, DepMap data revealed that peripheral nervous system (PNS) tumors, including neuroblastomas or malignant peripheral nerves sheath tumors (MPNSTs), are sensitive to WSB2 depletion. Neuroblastomas are derived from primitive cells of the sympathetic nervous system and represent 15% of cancer death in children, with 90% of them diagnosed before the age of 5^68–72^. In aggressive neuroblastoma, MYCN deregulation correlates with an upregulation of the p53 negative regulator MDM2 as well as the anti-apoptotic factor BCL2^72^. Overactivation of the MAPK/PI3K/AKT pathways by BDNF factor deregulation leads to a suppression of BIM induction and drug resistant tumors^72^. As such, our finding suggests that CUL5^WSB2^ inhibitors may adversely affect growth of Neuroblastomas irrespective of the BCL2 family member by causing strong up-regulation of BIM. However, although our mechanistic studies have identified BIM as a major substrate of WSB2 in RPE1, we have not directly demonstrated that the apoptotic phenotype observed in neuroblastoma cells is primarily mediated by BIM, as other WSB2 substrates could contribute to this phenotype. Firmly establishing WSB2-BIM relationship with apoptosis in neuroblastoma will require additional work and remains an important question.

In this study, we identified the CUL5^WSB2-BCL-XL/MCL1^ complex as a novel co-receptor for the regulation and degradation of BIM. Taking advantage of the powerful structure-based AlphaFold A.I^55^, we identified the interaction motif required for WSB2/BCL-XL binding, which is essential for BIM targeting. While the K/R-E-M motif in BCL-XL is shared by BCL2 and BCL-W, it is not seen in MCL1, and mutations in WSB2 that disrupt BCL-XL/BCL2/BCL-W binding don’t affect MCL1 binding. Thus, unlike many other E3s which recognize a common degron in a disparate set of substrates, WSB2 recognizes functionally related substrates through independent mechanisms. Our results clearly indicate that the formation of the BCL-XL-BIM or MCL1-BIM dimers are necessary for WSB2 interaction. We do not yet understand why WSB2 fails to bind BCL-XL alone, as AlphaFold2 predicts this interaction to occur. BCL-XL is known to undergo conformational changes in response to interactions with pro-apoptotic proteins, and these may be required for binding^73^.

## Supplemental Information

**Figure S1:**
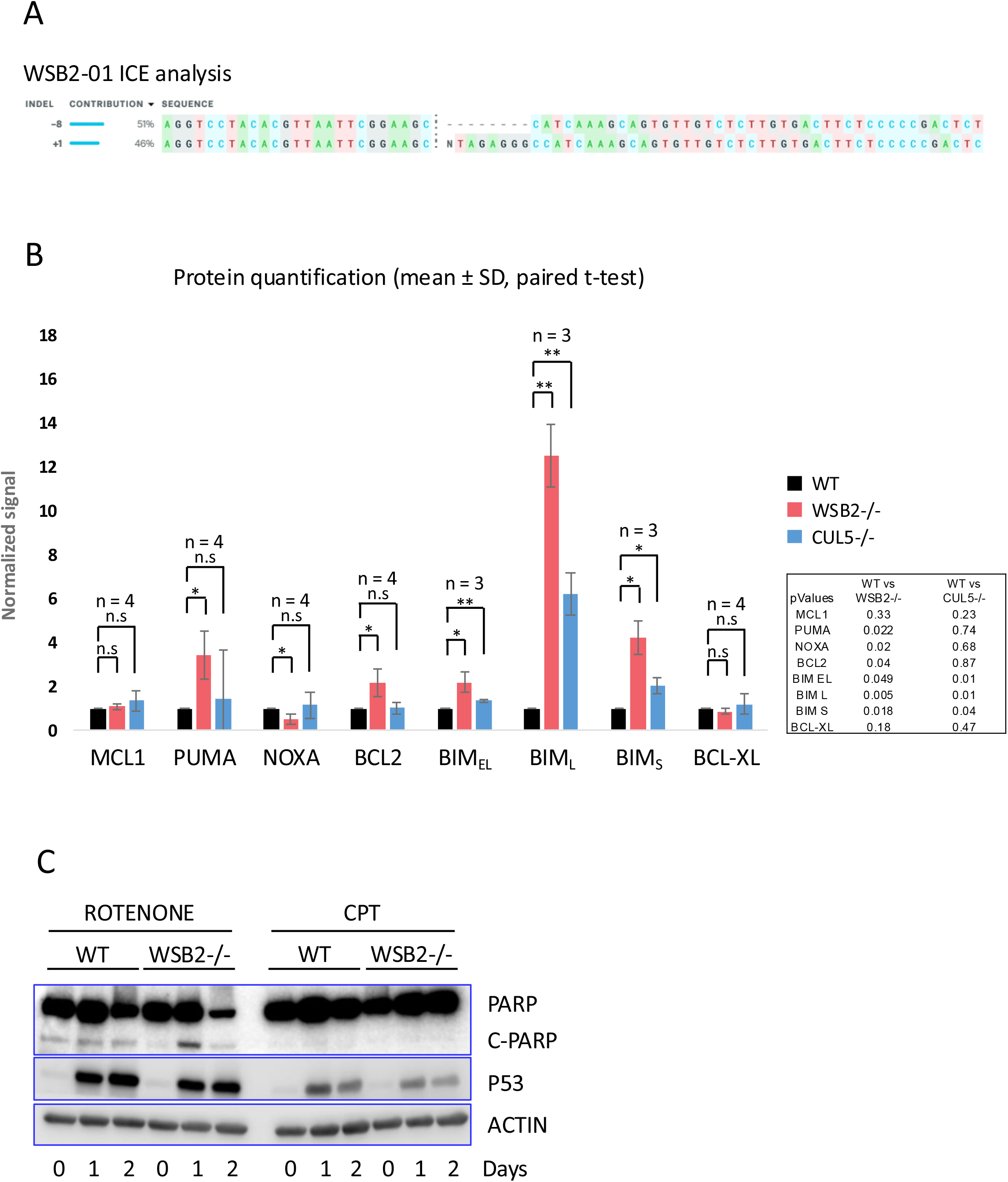
WSB2 clone sequencing validation and sensitivity to mitochondrial stressors. **A.** ICE analysis results of Sanger sequencing results from the WSB2 CRISPR clone generated for this study. **B.** Figure 1D Immunoblot quantification of MCL1, PUMA, NOXA, BCL2, BIM_EL_, BIM_L_, BIM_S_ and BCL-XL relative protein signal level to RPE1 wild-type of WSB2^−/−^ and CUL5^−/−^ mutants cell extract. **C.** Immunoblot of wild-type and WSB2 mutant cell extract following treatment with Rotenone or Camptothecine for 48h.

**Figure S2:**
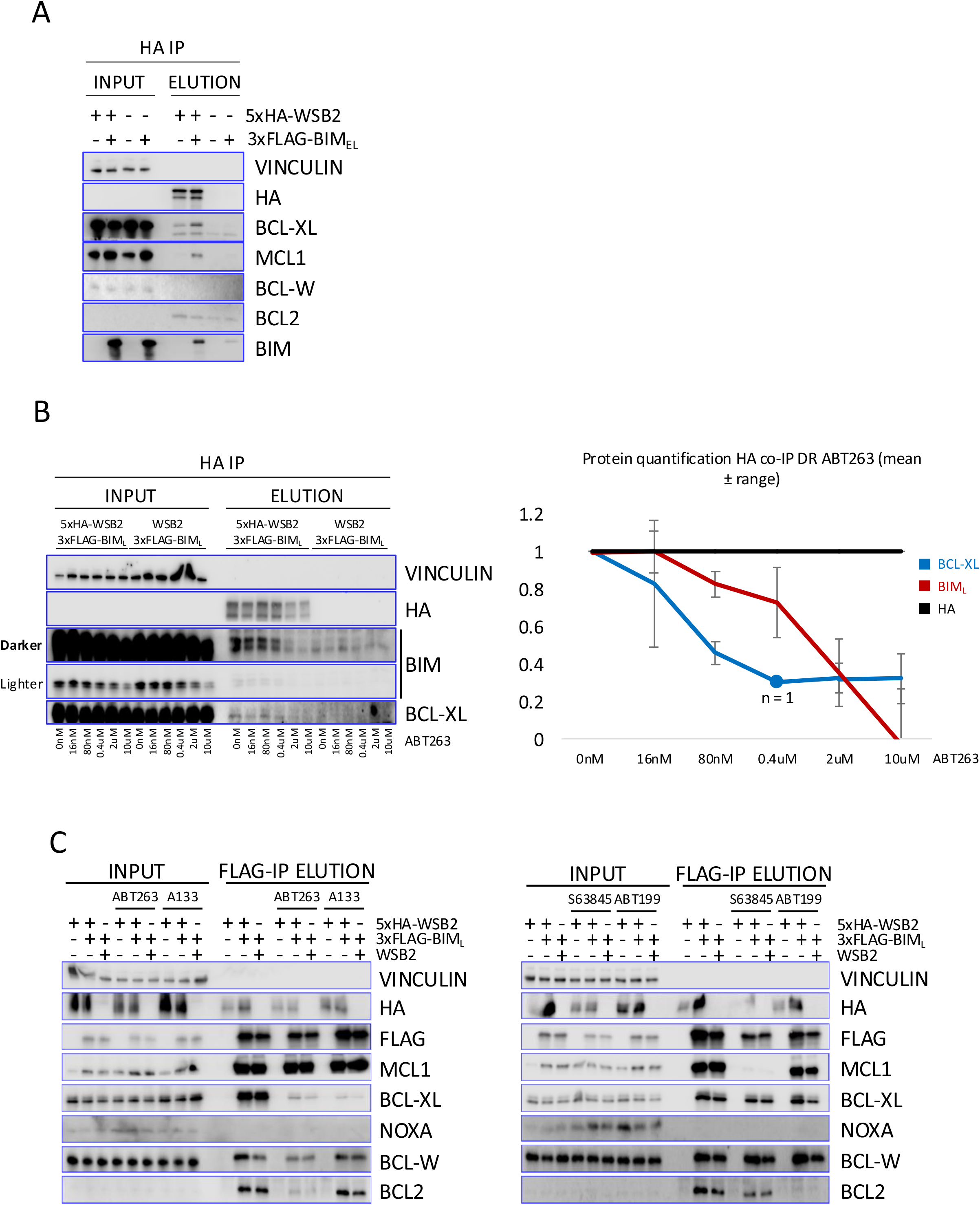
WSB2 interaction with endogenous BCL-W and BCL2. **A.** As figure 2C, except cell extract was blotted for BCL2 and BCL-W. **B. (Left)** HA co-immunoprecipitation immunoblot. Cells expressing a 3xFLAG-tagged BIM_L_ allele with either an untagged or 5xHA-tagged WSB2 allele were treated with increasing dose of ABT-263 3 hours prior to immunoprecipitation. **(Right)** Quantification of BCL-XL and BIM relative to the amount of HA-tagged WSB2 pulled down. The 0.4 uM BCL-XL point has only one repeat. **C.** Co-immunoblot analysis of 3xFLAG-BIM_L_ in the presence of the BH3 mimetics ABT-263, ABT-199, S63845, or A1331852. Immunoprecipitation of 3xFLAG-BIM was used to confirm the efficacy of each BH3 mimetic in disrupting its interaction with the respective anti-apoptotic proteins.

**Figure S3:**
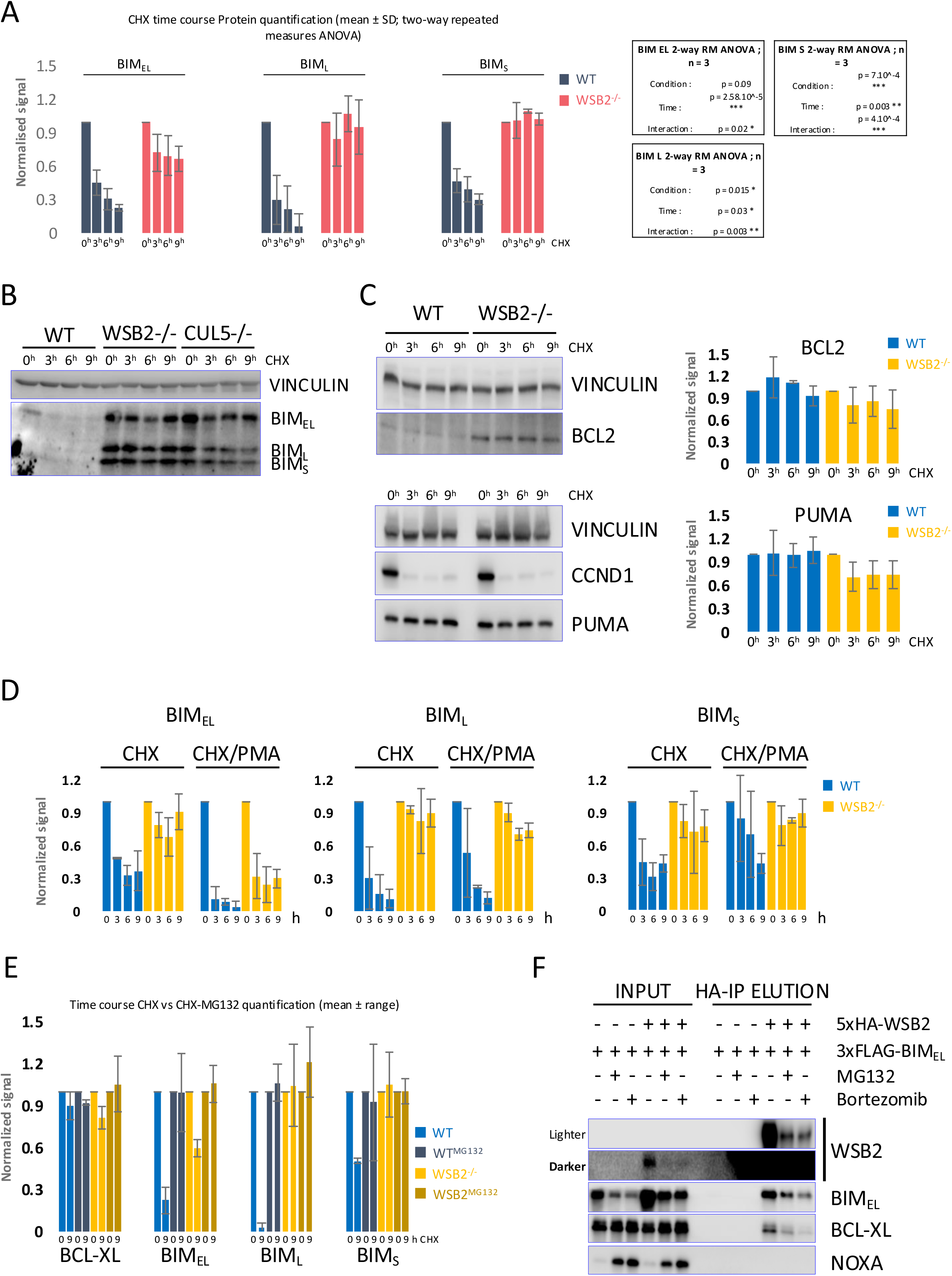
Quantification of CHX/PMA treatment. **A.** Quantification of each BIM isoform in wild-type and WSB2 mutant RPE1 cells, following treatment with cycloheximide, from figure 3A immunoblots. Signals were normalized to vinculin loading control and subsequently normalized to the t=0 signal. **B.** Immunoblot of BIM in wild-type, WSB2 and CUL5 mutant RPE1 cells, following a 9h time course with cycloheximide. **C. (Left)** Immunoblot of BCL2 and PUMA in WT and WSB2 mutant RPE1 cells after treatment with cycloheximide. Samples were harvested as indicated. **(Right)** Quantitation of BCL2 and PUMA levels were normalized to VINCULIN and subsequently normalized to the t=0 signal. **D.** Quantification of figure 3B immunoblots in WT and WSB2 mutant RPE1 cells with or without PMA added to one set of samples at the same time as cycloheximide. The quantifications were performed as in figure S3A. Phosphoforms of each BIM isoforms were combined for normalization. **E.** Quantification of figure 3C immunoblots in WT and WSB2 mutant RPE1 cells with or without MG132 added to one set of samples at the same time as cycloheximide. The quantifications were performed as in figure S3A. **F.** Western blot analysis of Bortezomib and MG132 effects on 5xHA-WSB2 and 3xFLAG-BIM_EL_ expression compared to untreated.

**Figure S4:**
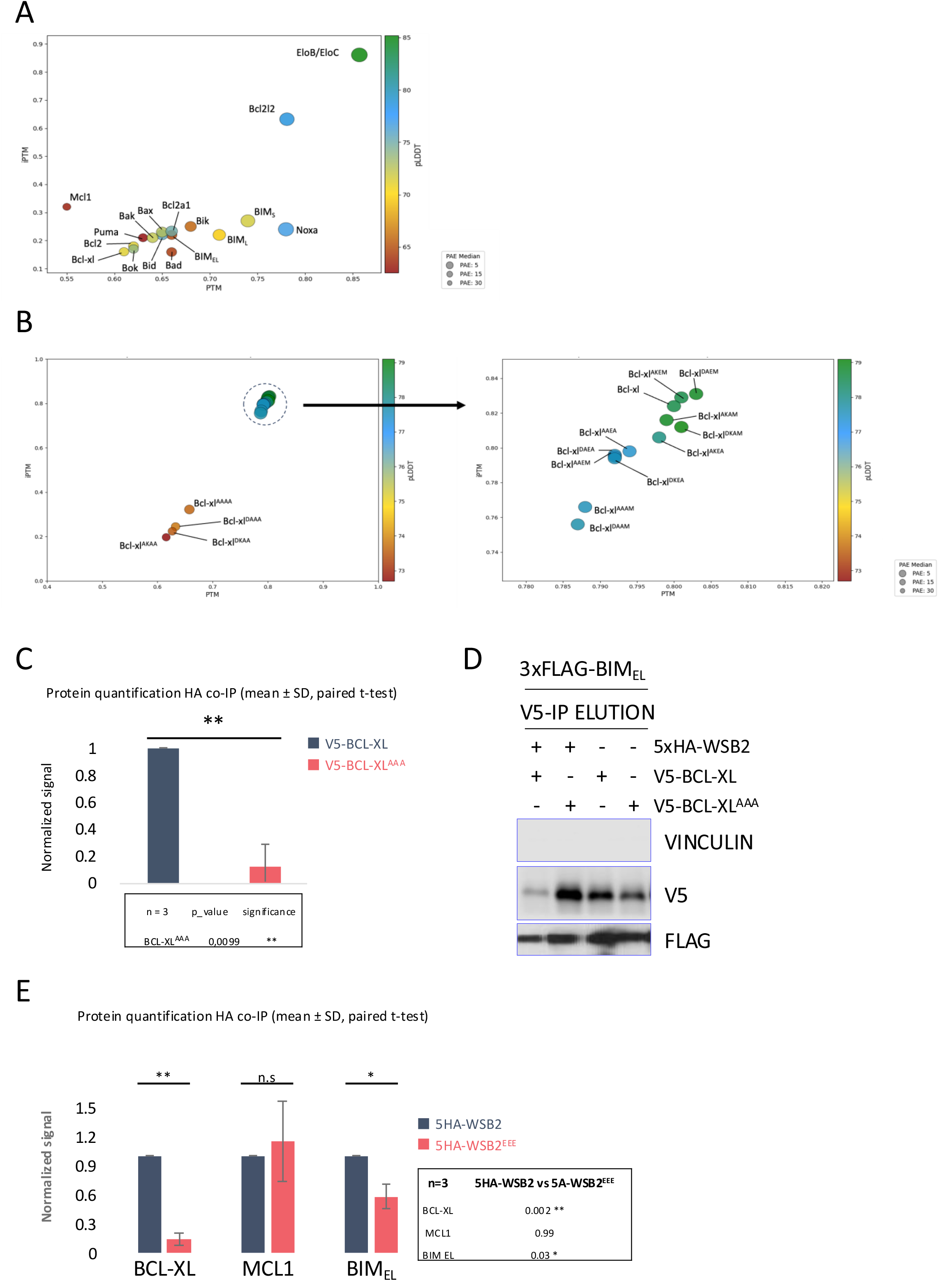
AlphaFold2 MM plots. **A.** AlphaFold 2 multimer predicted WSB1 dimerization with BCL2 protein family members scores. The CUL5 substrate adaptors ELOB and ELOC were used as a positive control. **B.** AlphaFold 2 multimer predicted WSB2 dimerization with BCL-XL DKEM motif mutants scores. Right Panel represent a zoomed in window of the upper right cluster of predicted interaction in the left panel. **C.** HA-immunoprecipitation relative protein signal quantification of figure 4D. V5-tagged BCL-XL or V5-tagged BCL-XL^K157A/E158A/M159A^ allele (BCL-XL^AAA^) were expressed in RPE1 cells co-expressing a 5xHA-WSB2 allele. **D.** V5 immunoprecipitation experiment in cells expressing an untagged or 5xHA-tagged WSB2 allele co-expressed with either V5-tagged BCL-XL or BCL-XL^K157A/E158A/M159A^ (BCL-XL^AAA^). In all cell lines, a 3xFLAG-tagged BIM_EL_ allele was co-expressed. **E.** HA-immunoprecipitation relative protein signal quantification of figure 4F. Untagged WSB2, 5xHA-tagged WSB2^R157/R174/R297^ or wild-type WSB2 were co-expressed with 3xFLAG-BIMEl in RPE1 cells.

**Figure S5:**
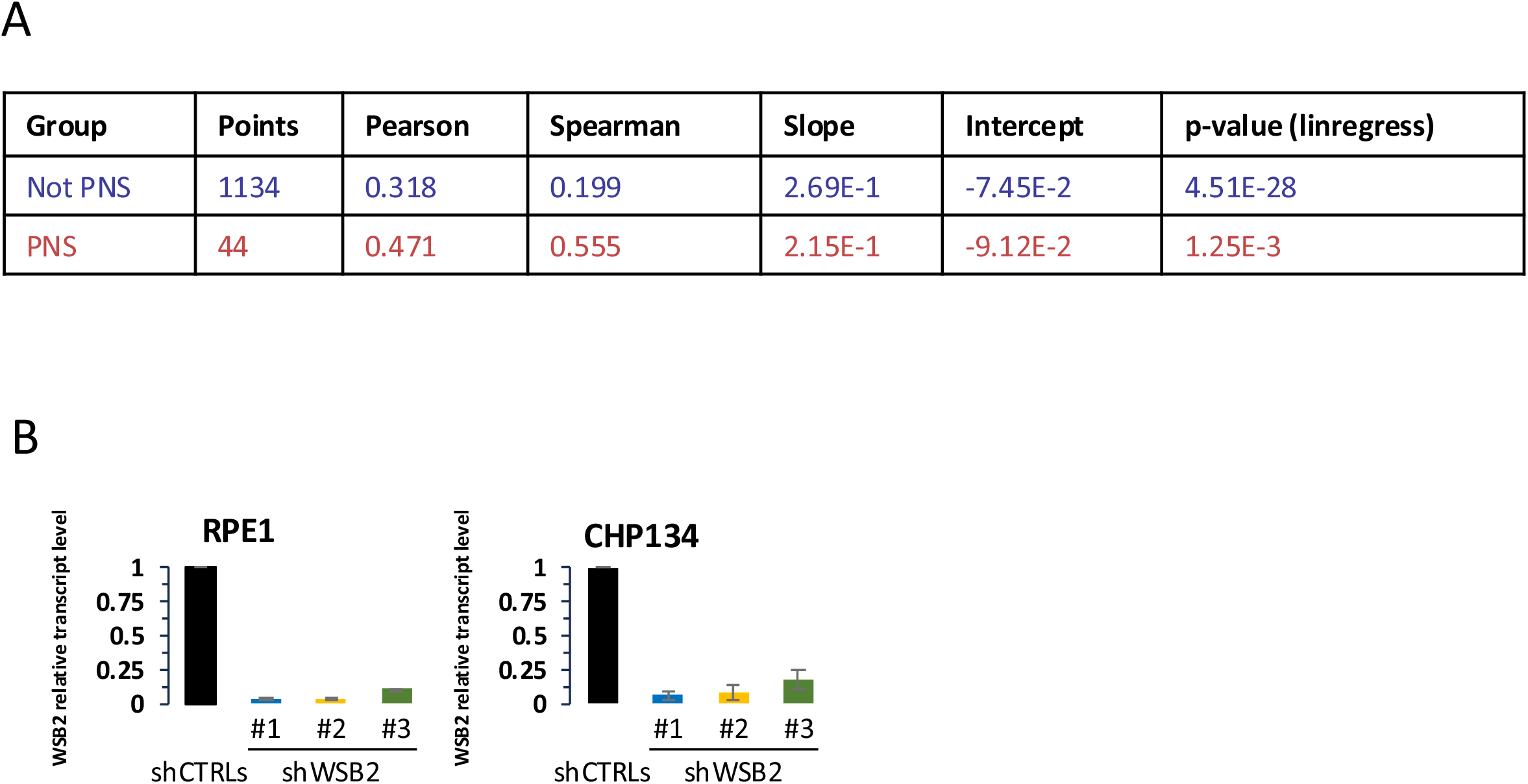
DepMap statistics & shRNA controls. **A.** DepMap generated regression line for Peripheral Nervous System (PNS) tumors and all other tumors types (referred to as not PNS) for sensitivity to WSB2 versus BCL2L2 (BCL-W) depletion **B.** Quantitative PCR of WSB2 transcript levels. Constitutive expression of a WSB2 targeting shRNA in RPE1 or CHP134 cells for 24 hours decrease WSB2 transcript level compared to scramble. Data are represented as mean ± SD

## Materials & Methods

### Cell Culture

#### Cell Lines

Cell lines were maintained using standard tissue culture techniques. HEK293T and RPE1 were cultured in DMEM F/12 (Thermofisher #11320082) with 10% heat inactivated Fetal Bovine Serum (Gibco #A52568-01) and 1X Anti-Anti (Gibco #15240-062). HAP1 were cultured in IMDM (Thermofisher # 12-440-061) with 10% heat inactivated Fetal Bovine Serum and 1X Anti-Anti. CHP134 were cultured in RPMI (Thermofisher #21870-076), 2mM L-glutamine (Sigma-Aldrich #SLCQ1268), 20% heat inactivated Fetal Bovine Serum and 1X Anti-Anti. TGW were cultured in EMEM (Corning #10-009-CV) with 20% heat inactivated Fetal Bovine Serum and 1X Anti-Anti. All cells grew at 37°C with 5% CO2.

#### Lentiviral particle generation & transduction

Lentiviral particles were produced using HEK293T seeded at ∼6×10^5 cells into a 6-well plate (∼30-40% confluency) and transfected the following day (∼80% confluency) with 1 µg of the vector of interest, 750 ng pSPAX2, 250 ng VSVG, and 6 µL Fugene® HD (Promega #E2311) in 100 µL Opti-MEM® (Gibco #31985-070). The transfection mix was incubated 15 minutes at RT and added dropwise to cells. After 6 hours at 37°C, the medium was replaced with fresh media. Virus-containing supernatants were collected after 24 hours, filtered (0.45 µm, Thermo Scientific #723-2545), and stored at −80°C. Target cells were transduced with a 1:10 dilution of virus-containing media in the presence of 5 µg/mL Polybrene or 8 µg/mL Protamine Sulfate.

#### Cell lines generation

RPE1 clonal lines were generated by sequential lentiviral transduction with plasmids carrying dox-inducible CRISPR-Cas9 systems targeting CUL5 or WSB2 (gRNA sequences: CUL5 - ACTATCTTAGCTGAGTGCCA, WSB2 - CACGTTAATTCGGAAGCTAG; Toronto CRISPR Human Knockout Library, TKOv3). Cells were selected in 1 µg/mL or 10 µg/mL puromycin, induced with 1 µg/mL doxycycline for 3 days, and sorted into single-cell clones by FACS. Clones were validated by western blotting (CUL5) or sequencing (WSB2) (Fig. S1A). Overexpressing cell lines were obtained by transfecting cells with either the pH198 or pTB97 plasmid carrying a TRE3GS promoter and Tet-ON system. The WSB2/5xHA-WSB2 alleles were transfected on the same plasmid with a PuromycinR selection marker, BIM alleles were transfected with a GFP selection marker and BCL2/BCLXL/V5-BCLXL were transfected with a BlasticidinR selection marker. Cells were selected for their respective resistance and/or sorted by FACS for GFP positive signal.

## Method details

### Time Course Experiments

Drugs concentration used outside of sensitivity assay are 10uM ABT-263 Selleck Chemicals #S1001; 500nM A-1331852 Selleck Chemicals #S7801; 15uM S63845 TargetMol #T5346; 15uM ABT-199 TargetMol #T2119; cycloheximide 50uM Sigma-Aldrich #C4859-1ML; 100nM phorbol 12-myristate 13-acetate (PMA) MCE #HY-18739; 500 nM MLN4924 MedChem Express #HY-70062; 50-100uM Z-VAD-FMK AdooQ BioScience #A12373

### qPCR Analysis

Total RNA was extracted from 10⁵–10⁷ cells using TRIzol (ThermoFisher #15596018) according to the manufacturer’s protocol, purified, and DNase I-treated using the RNA Clean & Concentrator™-5 Kit (Zymo Research #R1015). RNA purity was confirmed by nanodrop quantification and/or agarose gel electrophoresis. 5ug of RNA were reverse-transcribed using the Tetro RT-PCR Kit (Meridian Bioscience #BIO-65043). Primers (75–150 bp, exon-exon junctions when possible) were designed using NCBI Primer-BLAST and validated with standard curves (5×10-fold dilutions). Relative expression levels were normalized using the ΔΔCt method with Actin as the housekeeping gene. Primer sequences are:

- **CUL5**: Forward: GCAAGCACAGGCACGAGT; Reverse: TTGCTGCCCTGTTTACCCAT
- **WSB2**: Forward: CTGAGGTCACTCCACCACAC; Reverse: GGAGTCTGTCATCTGCCACC
- **BIM**: Forward: ATGTCTGACTCTGACTCTCG; Reverse: CCTTGTGGCTCTGTCTGTAG
- **NOXA**: Forward: AGAGCTGGAAGTCGAGTGTG; Reverse: GAAACGTGCACCTCCTGAGA
- **MCL1**: Forward: GTTTTCAGCGACGGCGTAAC; Reverse: CTCCACAAACCCATCCCAGC
- **BCL-XL**: Forward: TCAATGGCAACCCATCCTGG; Reverse: GAGCTGGGATGTCAGGTCAC
- **ACTIN**: Forward: AGGCACCAGGGGGTGAT; Reverse: GCCCACATAGGAATCCTTCTGAC

### Western Blotting

#### Protein extraction

Proteins were extracted by resuspension and mechanical lysis in lysis buffer (50% 2X RIPA buffer (100mM Tris pH8; 300mM NaCl; 2% TritonX100; 1% Sodium Deoxycholate; 0,2% SDS); 80mM Beta-Glycerophosphate; 5mM NaF; PepA; Protein Inhibitor cocktail; 2mM Sodium orthovanadate; 1mM PMSF or 50% 2X RIPA buffer; cOmplete Tablets, Mini EDTA-free, *EASYpack* (Roche #04693159001) and PhosSTOP *EASYpack* (Roche #04906837001) following manufacturer recommendation). After 30 minutes on ice, lysates were clarified by centrifugation at 21,000 × g for 15 minutes at 4 °C. Protein concentrations were determined via BCA assay, then normalized lysates were mixed with Laemmli buffer (50 mM DTT) and heated at 95 °C for 5 minutes.

#### SDS-PAGE and transfer

Proteins were loaded at equal quantity on Criterion 26- or 18-wells TGX 4-20% Precast polyacrylamide gel (BIORAD #5671095; #5671094) and ran at constant Amperage in Running buffer (384mM Glycine; 50mM Tris-Base; 0.1% SDS(w/v)). The proteins were subsequently transferred to a 0.2uM Nitrocellulose blotting membrane (Amersham Protan #10600001) at constant amperage; 100V for 1h at 4°C using a BIORAD PowerPack HC, in transfer buffer (20mM Tris-Base; 150mM Glycine; 0,0374% SDS (w/v); 2% Methanol (v/v)). Membranes were subsequently blocked in PBS-T (0.05% Tween) 5% milk for 1h at RT and incubated overnight with primary antibodies.

#### Detection method

Membranes were washed 3 times in PBS-T (0.05% Tween) for 5 minutes each time before being incubated with secondary HRP antibody for 1h at RT in PBS-T (0.05% tween) 5% Milk. Membranes were subsequently washed 3 times in PBS-T for 5 minutes each time. Proteins were revealed by incubating the membrane with regular or strong ECL (PROMETHEUS #20-300B or ThermoScientific #34096), depending on the signal quality, for 2 minutes at RT before imaging protein signal on a LICOR Odyssey. Protein signal quantification was performed using the ImageStudio LICOR software and Microsoft Excel.

### Immunoprecipitation/Co-immunoprecipitation

Cells were resuspended in lysis buffer (150 mM NaCl, 50 mM Tris pH 8, 0.5% Triton X-100) supplemented with PhosStop™ phosphatase and cOmplete™ protease inhibitors. Lysates were prepared by passing them 10 times through a 21-gauge needle, then incubated for 30 minutes at 4 °C with rotation and cleared by centrifugation at 21,000 × g at 4°C for 20 minutes. Protein concentrations were measured by BCA assay. For HA-tag IP, lysates were incubated 30 minutes at RT with anti-HA–coated beads (Pierce or Dynabeads Protein A preloaded with anti-HA). For FLAG-tag IP, lysates were incubated 30 minutes (or 1 hour for ubiquitin IP) with anti-DYKDDDDK beads. For V5-tag IP, lysates were pre-incubated with anti-V5 antibodies, then with Dynabeads. For MYC-tag IP, lysates were incubated 1 hour at RT with anti-c-MYC beads and washed in 50% 2× RIPA. Proteins were eluted at 65 °C for 10 minutes, reduced in Laemmli buffer with 50 mM DTT, and heated to 95 °C for 5 minutes before western blotting. A WSB2 untagged allele was used as negative control in HA pull-down. In the WSB2/BCL2 co-IP experiment no allele of WSB2 was overexpressed in the negative control.

### AlphaFold Analysis

Protein FASTA sequences were obtained on the Uniprot database. Default settings were used for collabfold version 1.5.5. To evaluate the accuracy of our protein complex model predictions, we used the four parameters used by AlphaFold2 for predicted model confidence, PTM, iPTM, pLDDT, PAE. We considered iPTM > 0.8; PTM > 0.5; PAE < 10 and pLDDT > 77 to be significant.

### Internally controlled, competitive growth assays

shRNA constructs targeting WSB2 were ordered as pre-designed (Sigma-Aldrich) or synthesized (IDT) and cloned into the pLKO.1 vector. shRNAs lentiviral constructs (GFP^+^) were used to transfect cells, which were then sorted by FACS (SonySH800). The GFP^+^ fraction was mixed 1:1 with an untransfected GFP^−^ population in RPMI (20% FBS, 1x Anti-Anti, 2 mM L-Glutamine) or EMEM (20% FBS, 1x Anti-Anti) at T0. Each day, 3,500–10,000 cells per condition were resorted to quantify the GFP^+^/GFP^−^ ratio using a 99th-percentile gating strategy. The resulting ratios were normalized to the T0 value. The mean of each experiment was pooled and represented as bar graphs ± SD. The shRNAs sequences are: (sh1) CACGGCTTCTTACGATACCAA; (sh2) CCCTTCGAAGTTTCCTAACAA; (sh3) CCCACCAGTTTGATTGGAAGT; (shCTRL1) CAACAAGATGAAGAGCACCAA; (shCTRL2) CCTAAGGTTAAGTCGCCCTCG.

### Caspase-Glo Assay

At the starting point of the experiment (T0), 1200 viable, singlets wild-type (pre-infection) CHP134 cells were sorted by FACS and serially diluted in a 3-points 10-fold dilution series in a final volume of 25ul per well in a white-walled 96 multiwell Corning® Microplates (Cat.# 3917). According to manufacturer instructions, an equal volume of Caspase-Glo® reagent was added to each well, followed by a 30 minutes incubation at RT on a rotator, protected from light. Luminescent signal was subsequently measured using a Synergy II plate reader. After transfection, 1200 viable, singlets GFP+ CHP134 cells were sorted by FACS for each shRNA and wild-type control and processed identically. We used 1000 cells for our experiments, as such the luminescent signal was normalized to the number of cells and then to the wild-type control (used for that particular day) to assess apoptosis level. The mean of each experiment were pooled and represented as bar graphs ± Range.

### Antibodies for Western Blotting

1:500 CUL5 (#sc-373822), 1:1000 ELOC (#sc-135895), 1:1000 PUMA (#4976S), 1:1000 NOXA (#14766S), 1:1000 BIK (#4592T), 1:1000 BAD (#9292S), 1:1000 BCL2 (#4223S), 1:1000 BCL-w (#2724S), 1:1000 BCL-XL (#2764S), 1:1000 MCL1 (#94296S), 1:1000 BID (#2002T), 1:1000 BAK (#12105S), 1:1000 BAX (#5023S), 1:1000 BIM (#2933S), 1:1000 p53 (#9282S), 1:1000 HA (#3724S), 1:1000 FLAG (#14793S), 1:1000 MYC (#2276S), 1:1000 V5 (#13202S), 1:1000 MARCH5 (#v06-1036), 1:500 Cyclin D1 (#ab16663), 1:3000 ACTIN (#A5441-100UL), 1:3000 VINCULIN (#V9131).

## Acknowledgments

We thank the UCSF Helen Diller Family Comprehensive Cancer Center LCA and LCA-Genomic Core Facility for the service they provided in support of this research. We extend our thanks to Emma Bolech and Candace S.Y Chan for their contributions to this project. We also appreciate the Weiss laboratory for providing the Neuroblastoma CHP134 cell line with special thanks to Hiroyuki Yoda for his assistance. This work was supported by NIH grants R35GM118104, the UCSF Resource Allocation Program (RAP), the UCSF Program for Breakthrough Biomedical Research (funded in part by the Sandler Foundation) and the Cancer League.

## Author Contributions

DPT: Conceptualization, Methodology, Writing – original draft, Funding acquisition, Supervision, Resources, Project administration, Writing – review & editing

WVZ: Conceptualization, Methodology, Writing – original draft, Investigation, Formal analysis, Validation, Data curation, Visualization, Writing – review & editing

EMCA: Investigation, Validation NG: Investigation

## Competing Interest Statement

The authors declare no financial or non-financial competing interests in relation to this work

## Notes

### Competing Interest Statement

The authors have declared no competing interest.

### Summary of Updates

This revised manuscript includes new experiments and analyses addressing the regulation of BIM by WSB2. We added quantification and statistical analysis to several figures, as well as proteasome inhibition experiments supporting proteasome dependent turnover of BIM isoforms. A new dose response experiment examines BH3 mimetics disrupts the WSB2 and BIM association in a dose response fashion. We also added experiments showing that WSB2 associates with overexpressed BCL2 and that mutation of WSB2 proposed binding interface reduces this association (Same as BCLXL). The figures, supplementary data, methods, and discussion have been updated accordingly. We have also clarified the limits of the evidence concerning direct protein binding and the cause of apoptosis following WSB2 depletion.

